# GSEA-InContext: Identifying novel and common patterns in expression experiments

**DOI:** 10.1101/259440

**Authors:** Rani K. Powers, Andrew Goodspeed, Harrison Pielke-Lombardo, Aik-Choon Tan, James C. Costello

**Affiliations:** Computational Bioscience Program, University of Colorado Anschutz Medical Campus, Aurora, CO, 80045, USA.; Department of Pharmacology, University of Colorado Anschutz Medical Campus, Aurora, CO, 80045, USA.; Department of Medical Oncology, University of Colorado Anschutz Medical Campus, Aurora, CO, 80045, USA.

## Abstract

**Motivation:** Gene Set Enrichment Analysis (GSEA) is routinely used to analyze and interpret coordinate changes in transcriptomics experiments. For an experiment where less than seven samples per condition are compared, GSEA employs a competitive null hypothesis to test significance. A gene set enrichment score is tested against a null distribution of enrichment scores generated from permuted gene sets, where genes are randomly selected from the input experiment. Looking across a variety of biological conditions, however, genes are not randomly distributed with many showing consistent patterns of up- or down-regulation. As a result, common patterns of positively and negatively enriched gene sets are observed across experiments. Placing a single experiment into the context of a relevant set of background experiments allows us to identify both the common and experiment-specific patterns of gene set enrichment.

**Results:** We compiled a compendium of 442 small molecule transcriptomic experiments and used GSEA to characterize common patterns of positively and negatively enriched gene sets. To identify experiment-specific gene set enrichment, we developed the GSEA-InContext method that accounts for gene expression patterns within a user-defined background set of experiments to identify statistically significantly enriched gene sets. We evaluated GSEA-InContext on experiments using small molecules with known targets and show that it successfully prioritizes gene sets that are specific to each experiment, thus providing valuable insights that complement standard GSEA analysis.

**Availability and Implementation:** GSEA-InContext is implemented in Python. Code, the background expression compendium, and results are available at: https://github.com/CostelloLab/GSEA-InContext

## 1 Introduction

Gene Set Enrichment Analysis (GSEA) [42, 37] was developed to help with the analysis and interpretation of the long lists of genes produced from high-throughput transcriptomic experiments. By summarizing genome-wide gene expression changes into gene sets - sets of functionally related genes - a user can gain insight into how biological pathways and processes are affected under the tested experimental conditions. Since its initial application to microarray experiments, GSEA has demonstrated utility across many applications including RNA-seq gene expression experiments, genome-wide associations studies [53, 13], proteomics [32], and metabolomics studies [50].

The power of GSEA lies in its use of gene sets, which provide a more stable and interpretable measure of biological functions compared to individual genes that can show greater experimental and technical variation [16, 49]. A user can define custom gene sets, but more commonly, researchers rely on pre-compiled gene sets, such as the widely-used Molecular Signatures Database (MSigDB) [42]. Additional online resources have become available to provide pre-compiled gene sets specific to drug response [52], human disease and pharmacology [1], molecular phenotypes [26], and patient prognosis [11], to name a few.

Similar to other Functional Class Scoring (FCS) methods [29], the underlying hypothesis of GSEA is that genes involved in a similar biological process or pathway (grouped into gene sets) are coordinately regulated. Thus, if an experimental perturbation activates a pathway, the genes in this gene set will be coordinately up-regulated and this pattern can be identified using statistical tests. The enrichment score, which reflects the degree to which genes in a gene set are over-represented at either end of a ranked gene list, is a fundamental aspect of FCS methods. Accordingly, a great deal of effort has been devoted to the development and evaluation of statistical models, from simple mean/median gene level statistics [27] or maxmean statistics [15] to the Kolmogorov-Smirnov [42, 37] and Wilcoxon rank sum tests [5]. GSEA uses a modified version of the Kolmogorov-Smirnov statistic to compute the gene set enrichment score [42, 37]. Finally, the significance of the enrichment score is estimated against the null hypothesis. Two categories of null hypotheses are used across FCS methods: *i*) self-contained or *ii*) competitive null hypothesis. When running GSEA [42, 37], these options can be found under the “Permutation type” field with options, `phenotype` (self-contained) or `gene_set` (competitive).

The self-contained null hypothesis states that *no genes in a given gene set are differentially expressed*. To test this hypothesis for any given gene set, the phenotype labels defining the experimental condition of individual samples are permuted. This approach focuses on the genes in a given gene set and ignores genes outside the set, providing strong statistical power and rejecting more null hypotheses [29, 19, 44]. However, this approach has several drawbacks. For experiments with a high number of differentially expressed genes, this approach will produce many significantly enriched gene sets. Conversely, if few genes are differentially expressed, correspondingly few to no gene sets will be significantly enriched. Additionally, because phenotype labels are permuted under this null hypothesis, the statistical power of the test is determined by the number of samples in the experiment. As a result, the GSEA documentation recommends providing at least seven samples per phenotype label when running GSEA with the `phenotype` option selected in the “Permutation type” field [21]. Experiments with fewer than three samples per phenotype cannot be run, and tens to hundreds of samples per experimental condition are needed to achieve robust statistics.

For the large number of experiments generating less than seven samples per condition, the alternative to the self-contained null hypothesis is the competitive null hypothesis. The null hypothesis for this approach states that *genes in a given gene set are at most as often differentially expressed as the genes not in the set*. To test this, random sets of genes of equal size to a given gene set are scored. Thus, this approach compares genes within a set to genes outside the set. When sample sizes are numerous and the data follow the assumptions of the underlying statistical models, then the self-contained null hypothesis is preferred as it offers greater statistical power than the competitive null hypothesis to reject the null hypothesis [29, 19, 44]. However, when these assumptions are not met or the focus of an analysis is on an individual sample, the competitive hypothesis is needed. When running GSEA [42, 37], the competitive hypothesis can be selected using the `gene_set` option under the “Permutation type” field [21]. It is also the only option when running the “GSEAPreranked” mode, where the user supplies a pre-ranked list of genes based on whatever method they choose, most often this is a list of differentially expressed genes.

There are many experiments that require the use of the competitive null hypothesis for proper comparison. Accordingly, this requirement motivated a series of methods to address the statistical challenges in single-sample analysis of ranked gene lists [24, 33, 3, 45]. By selecting random sets of genes outside the set being tested, the competitive null hypothesis approach breaks the inherent correlation structure of genes in the tested set. Methods like GSVA [24] nicely address this challenge by incorporating gene-specific variation directly in the calculation of a sample-wise gene set enrichment score within a given input data set.

Here we take a different approach to analyze and adjust for patterns in differentially enriched gene sets produced using GSEA with the competitive null hypothesis. Specifically, we account for gene-specific variation estimated from an experimental background. Our approach is motivated by the fact that there are no methods available for a user to easily compare their GSEA results to GSEA results obtained in other experiments to discern similar and/or distinct patterns affected across experiments. Overall, the goal of this research is to address two questions: 1) which gene sets are commonly enriched across a compendium of experiments, and 2) which gene sets are uniquely enriched in a single experiment compared to many other, independent experiments?

To accomplish these goals, we first curated a compendium of gene expression experiments encompassing a variety of experimental conditions and identified patterns of positive and negative enrichment by applying GSEA. We then leverage these patterns to help contextual single experiments. Accordingly, we developed an extension for GSEA that uses these context-specific patterns to inform the statistical testing procedure. Specifically, while GSEA tests for the significance of an enrichment score against a null distribution of enrichment scores calculated for *random* permuted gene sets, our algorithm generates permuted gene lists based on a set of user-defined background experiments. Because we allow the user to define the context of the background set of experiments, we have termed our method, GSEA-InContext, which stands for GSEA – <u>I</u>dentifying <u>n</u>ovel and <u>Co</u>mmon pa<u>tt</u>erns in <u>ex</u>pression experimen<u>t</u>s.

We applied GSEA-InContext to a compendium of gene expression experiments testing small molecule treatments in human cell lines. Small molecules remain the gold standard of treatment for numerous diseases, and in the context of cancer, human cell lines have been widely used to study mechanisms of drug action and present a robust pharmacogenomic platform [20, 17, 41, 4]. Gene expression experiments are regularly performed to study the direct effect of a small molecule, but expression profiles will capture both on- and off-target effects of the small molecule and disentangling these effects remains a challenge. At the same time, patterns of positive or negative enrichment can provide insights into common (i.e. not tissue- or drug-related) responses to small molecule treatment. In this article, we demonstrate how GSEA-InContext can be used to gain insights into both aspects of small molecule treatment. We proceed by first describing our curated background compendium of small molecule gene expression experiments. We present an analysis of this compendium and identify commonalities in differentially expressed genes and significantly enriched gene sets, motivating the development of the GSEA-InContext method. Finally, we demonstrate GSEA-InContext on two example applications: Notch inhibition in T-cell acute lymphoblastic leukemia and investigating gene expression changes in response to dexamethasone and estradiol treatment in breast cancer cell lines.

## 2 Materials and Methods

### 2.1 Data collection and normalization

We queried the Gene Expression Omnibus (GEO) database [14] for human gene expression studies performed on the Affymetrix Human Genome U133 Plus 2.0 Array that tested small molecules. We excluded studies that had fewer than 2 replicates per condition, or that did not have an appropriate vehicle control condition, which was needed to calculate consistently controlled differentially expressed gene lists across all experiments. We proceeded with a total of 128 studies comprised of 2,812 individual microarrays that met the search criteria. Meta-data for each study was parsed from GEO in order to annotate tissue type, cell line, and small molecule. The CEL files for each study were downloaded with the *GEO*-*Query* R package [12]. Within each study, the expression data was background corrected, quantile normalized, and probe sets were summarized using RMA [6] with the *affy* R package [18]. For each study, control and treatment conditions were identified and differential expression between all control/treatment pairs was calculated with the *limma* package [39]. Probe sets were annotated to genes using the *hgu133plus2.db* R package [7], keeping the one probe set per gene with the highest average expression across all samples. For each experimental comparison, genes were ranked according to their log_2_ fold change and saved as a ranked list L (Figure 1A) for input into GSEAPreranked and GSEA-InContext. In total, we generated a compendium of 442 ranked lists.

**Figure 1:**
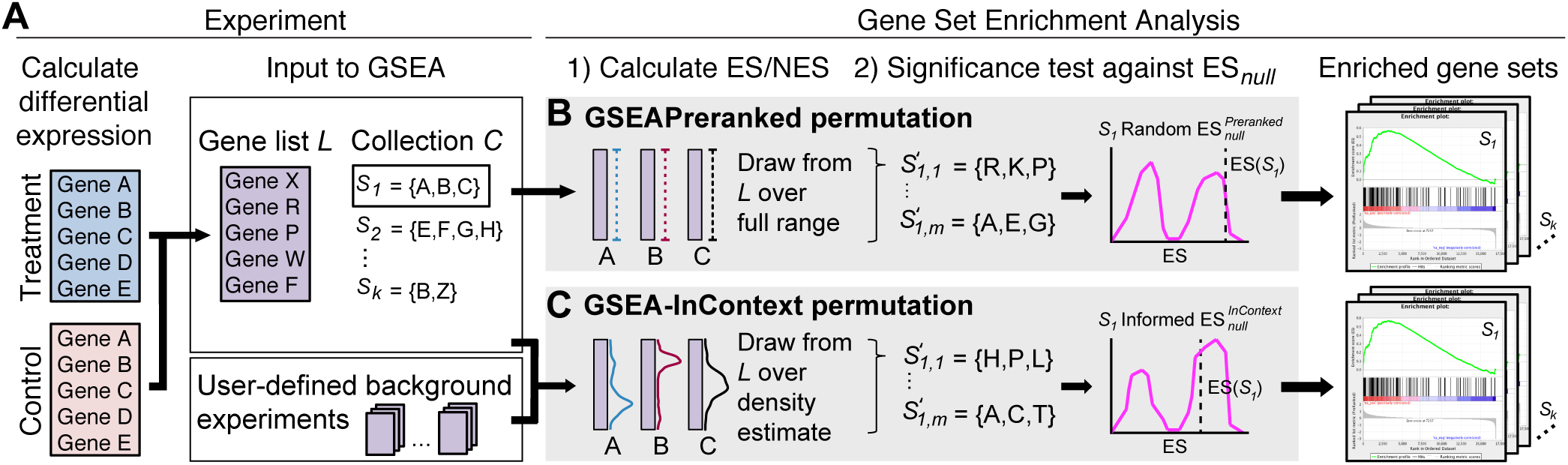
Overview of the statistical test for GSEAPreranked and GSEA-InContext. (A) A typical workflow for using GSEA to identify significantly enriched gene sets in a vehicle control vs drug-treated experiment. Calculating the expression fold change between the two conditions produces a ranked gene list L. This list is input into GSEA along with a collection of gene sets C. (B) To test whether a gene set *S*_1_ is significantly enriched in L, the enrichment score, *ES*(*S*_1_), is tested against a null distribution *ES_null_*. GSEAPreranked creates 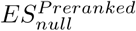 by calculating the *ES* for *m* gene sets the same size as *S*_1_, which are created by randomly selecting from teh full range of L. (C) The GSEA-InContext approach takes as input L, C, and a user-defined set of background experiments. Instead of randomly generating gene sets, 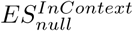 is created by selecting genes based on how they are distributed in the background set of experiments. The ES for each of *m* informed gene sets is calculated and used to evaluate the significance of *ES*(*S*_1_).

All gene set collections used were downloaded from MSigDB, v6.1 [42, 37, 34]. The Hallmarks collection [34] was selected to be used for all analyses because it is comprised of 50 gene sets, thus full results can be reported and displayed through this manuscript. Analyses performed with additional gene sets are supplied as described in Section 2.4.

To annotate mechanisms of action for the small molecules, we grouped them based on their targets using the Drug Repurposing Hub [10] and DSigDB [52].

### 2.2 Application of GSEAPreranked

To ensure consistency between implementations of GSEA, we ran each of the 442 ranked lists through the GSEAPreranked algorithm using both the javaGSEA Desktop program [42, 37] and the GSEApy Python package (https://github.com/BioNinja/gseapy); both implementations produced equivalent results. For all analyses shown here, we applied GSEApy (pypi package version 0.9.3, Python3.6) using a `weighted` enrichment scoring statistic and 100 permutations. GSEAPreranked requires the use of the competitive null hypothesis, the gene set permutation type. Default settings were used for all other parameters.

### 2.3 Implementation of GSEA-InContext

According to the GSEA documentation [21], the GSEAPreranked algorithm takes as input a user-supplied ranked gene list *L* and a collection of gene sets *C = {S_1_…S_k_}*, where *S_k_* is an *a priori* defined gene set (Figure 1A). An enrichment score *(ES)* is calculated for each gene set *ES(S*_k_) using a weighted Kolmogorov-Smirnov-like statistic [42, 37]. The *ES* reflects the degree to which genes in *S_k_* are positively or negatively enriched at either end of the ranked gene list *L.*

To estimate the significance level of *ES(S*_k_), GSEAPreranked tests *ES(S_k_)* against an empirically defined null distribution, 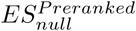. To illustrate how this distribution is created, we use the example of *S_1_* in Figure 1. GSEAPreranked generates *m* permuted gene sets of the same length as *S_1_* by randomly selecting genes from *L* (Figure 1B). We use the notation 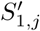 to represent the *jth* permutation of the randomized gene set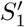. The nominal p-value for *S_1_* is calculated by comparing *ES(S_1_)* to the 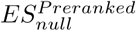 distribution. Note that the modified Kolmogorov-Smirnov test applied by GSEA creates a bimodal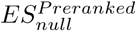.

Our method applies the same approach as GSEA to calculate the nominal p-value [42, 37, 21]. In contrast to GSEAPreranked, GSEA-InContext employs an alternative significance testing procedure to generate the null distribution, in which the *m* permuted gene sets are generated using the density of gene ranks estimated from a set of user-defined background experiments (Figure 1C). For a gene present in gene list L, let random variable *X* = {*x*_1_…*x*_n_} represent the set of gene ranks across all background experiments where *x*_i_ is the gene’s rank in the *i*^th^ background experiment. We estimate a gene’s probability density *Ĝ*(*X*) using a Gaussian kernel over the *n* experiments in the background set as follows:

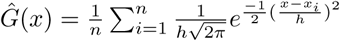

where *h* is the kernel bandwidth parameter. To enable the resolution of the kernel to scale with the size of the background set, we set the default for *h* to be the median distance between ranks across all observed ranks *x_1_…x_n_* for a gene. We also allow *h* to be set by the user. As shown in Figure 1C, *Ĝ(X)* is applied independently for all genes in a given gene set, such as *S_1_*, whose underlying density is to be estimated from the background experiments.

To create the permuted gene set 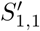, GSEA-InContext draws indices from *Ĝ(X)* for each gene in *S_1_*, then selects the gene at that index in *L.* This procedure is repeated to create m permutations of *S_1_* to generate the informed 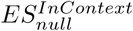. As in GSEAPreranked, the nominal p-value for *S_1_* is calculated by comparing *ES(S_1_)* to the 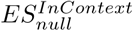 distribution (Figure 1C).

Outside of the changes to the way GSEA-InContext generates the null distribution of enrichment scores, all other components of GSEA are the same for GSEAPreranked and GSEA-InContext.

### 2.4 Code availability

To leverage the multi-threading capabilities of GSEApy, we implemented our method as a new class within the existing Python package. The background gene expression compendium of 442 ranked lists, the code to run GSEA-InContext, documentation, and supplemental results for all gene set collections are supplied at: https://github.com/CostelloLab/GSEA-InContext

## 3 Results

### 3.1 Overview of gene expression data sets

We curated a gene expression compendium of 442 gene lists ranked by log2 fold change between treatment versus control conditions. We required that all comparisons have at least 3 replicates per condition, where the conditions were either small molecule treatments or the appropriate vehicle control treatment. Raw data were processed according to the procedures outlined in Section 2.1. The tissues and small molecules included in the compendium are summarized in Figure 2. A total of 21 tissues are represented in the data set (202 unique immortalized or primary cell lines), with the most common tissue type being breast. We captured a range of 129 small molecules that we grouped into 69 drug classes based on mechanism of action. The most commonly used small molecules were eribulin and paclitaxel, which inhibit microtubule dynamics.

**Figure 2:**
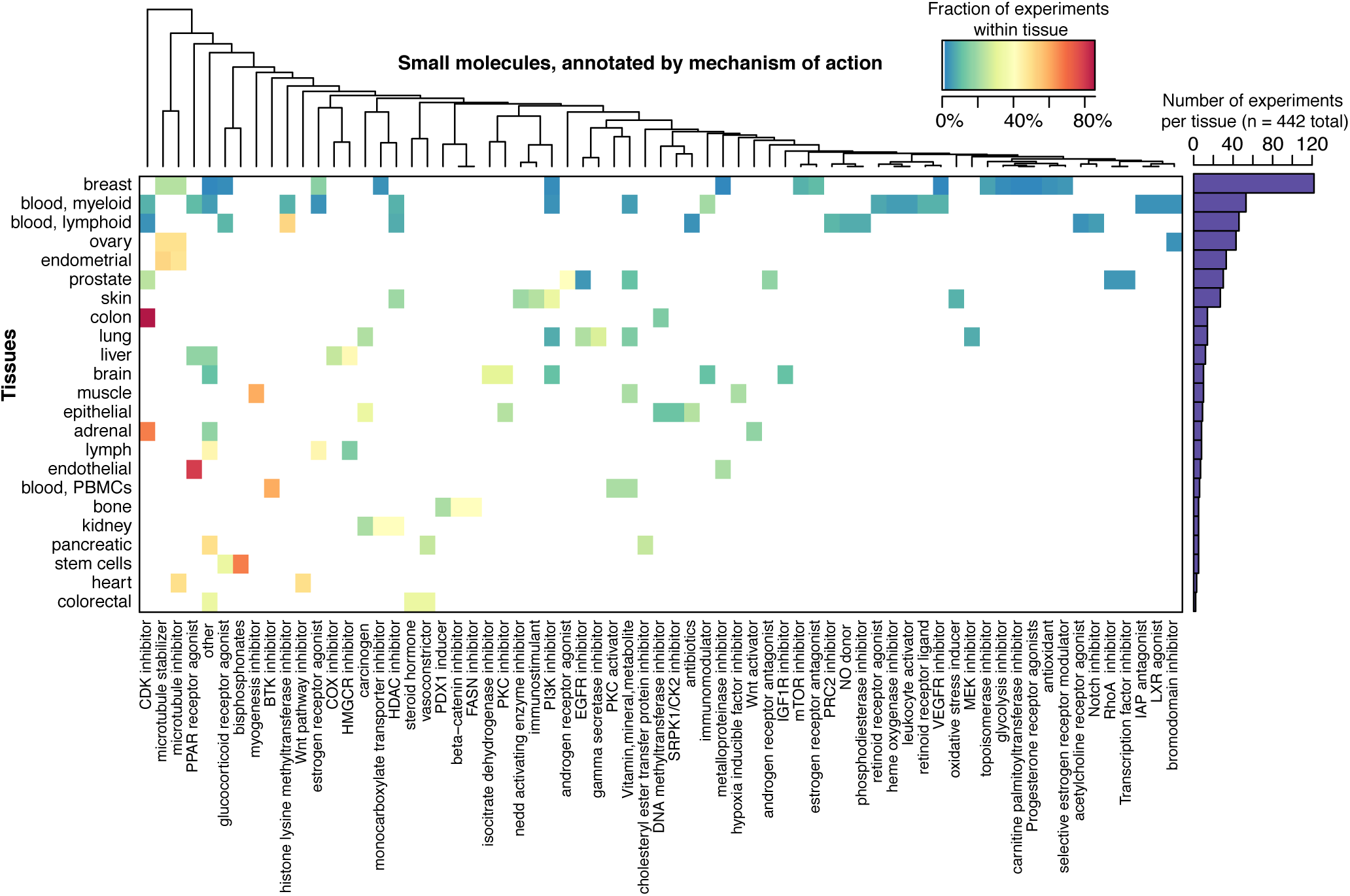
Overview of gene expression data sets by tissues and small molecules. Heatmap shows the fraction of small molecules used across 442 experiments (treatment vs. control comparisons). All experiments were performed in human immortalized or primary cell lines.

### 3.2 Common patterns of genes and pathways across small molecule treatments

To evaluate general gene- and pathway-level patterns, we first created a distribution of the mean rank for each gene across the 442 experiments. We compared these results to a null distribution generated by randomizing the genes in each of the 442 experiments. We found that roughly 25% of genes fell at least 3 standard deviations away from the mean rank of the null distribution, compared to the expected frequency of 0.3% (Figure 3A). Of the 25%, 12.6% of genes ranked higher and 13.9% ranked lower than the mean rank. These results demonstrate that roughly a quarter of the genes being studied across 442 experiments are more consistently differentially regulated than expected at random.

**Figure 3:**
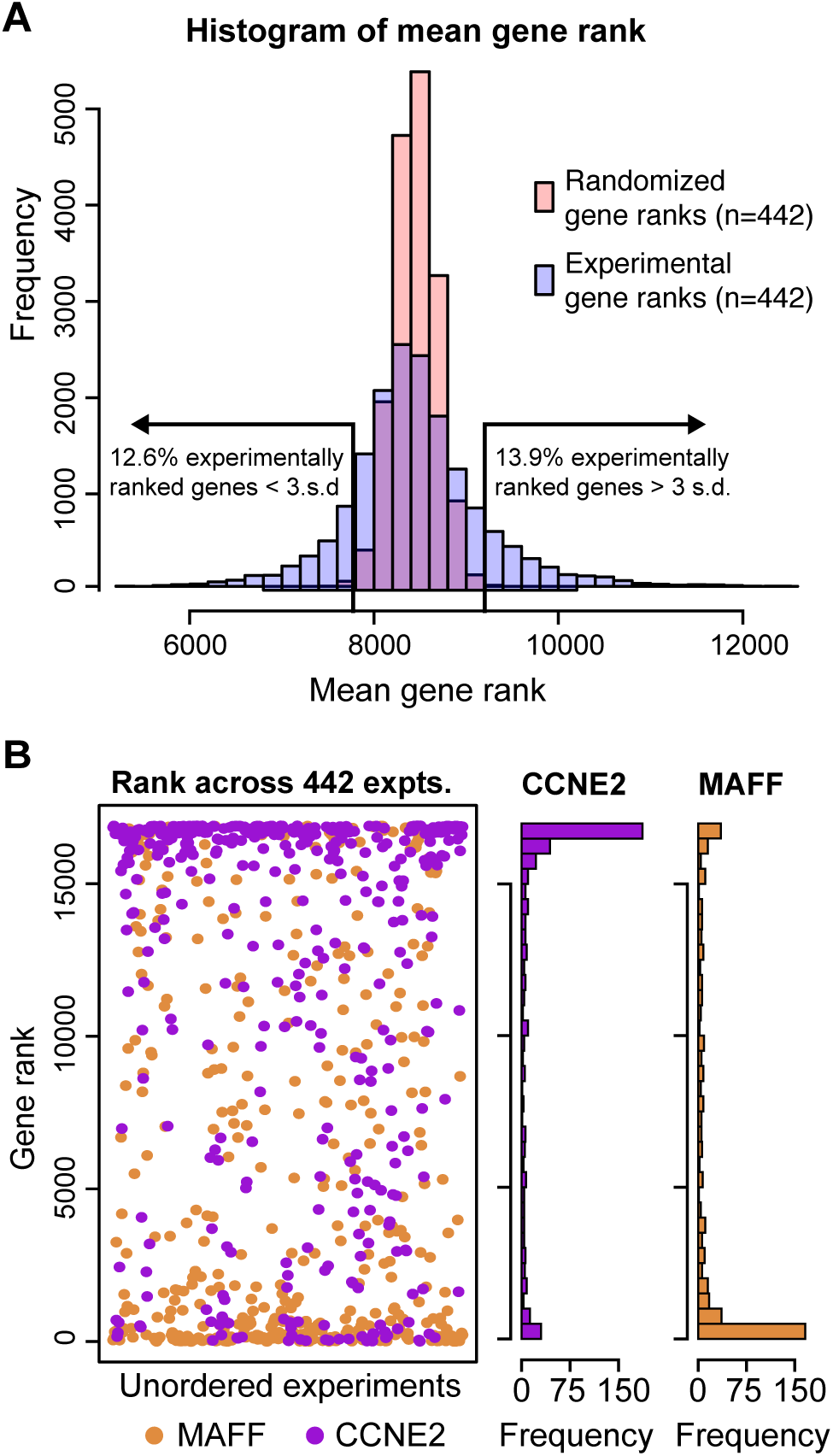
Ranking of genes across 442 small molecule gene expression profiles. (A) Distribution of the mean rank for all genes measured across 442 small molecule experiments (blue) compared to the mean rank of genes from 442 randomized gene lists (pink). Roughly 25% of genes in the experiments fall outside 3 standard deviations from the randomly ranked genes. (B) The ranks of MAFF and CCNE2 across all 442 experiments. These two genes are the highest and lowest ranked genes in (A) by mean rank across all 442 experiments.

To illustrate this effect on a per gene basis, Figure 3B displays the genes with the highest and lowest mean rank across all 442 experiments. The gene with the highest mean rank was MAF bZIP transcription factor F (MAFF), which encodes a transcription factor of the MAF family and has been shown to be essential for activation of genes involved in detoxification and the response to oxidative stress [28]. This gene is also up-regulated in response to hypoxia [9]. The most lowly ranked gene was cyclin E2 (CCNE2), an activating regulatory subunit of CDK2, most highly expressed during the G1/S cell cycle transition [22]. Intuitively, the rankings of these genes are consistent with small molecule treatment, given that CCNE2 is frequently down-regulated in response to drugs that arrest cell growth, and MAFF is up-regulated in response to cellular stress. However, the non-random ranking of genes does suggest there would be commonalities across enriched gene sets identified by GSEA. To investigate this, we ran GSEAPreranked on each of the 442 experiments and evaluated global gene set patterns using the Hallmarks collection [34]. We performed all analyses using all gene sets available in MSigDB [42, 37] and found similar patterns as those reported for the Hallmarks collection; these results are available as described in Section 2.4.

In Figure 4A, we report the fraction of experiments that showed an FDR < 0.05 for each of the gene sets in the Hallmarks collection, where we found clear patterns of positive and negative enrichment. For example, proliferation and cell cycle related processes were consistently down-regulated, including `E2_TARGETS`, which was significantly down-regulated in over 45% of the experiments. Other gene sets were consistently up-regulated, such as `TNFA_SIGNALING_VIA_NFKB`, which was signficantly positively enriched in approximately 53% of the experiments. These results are consistent with the trends we identified for CCNE2 and MAFF (Figure 3B). CCNE2 is a member of many of the down-regulated cell cycle-related gene sets and MAFF is a member of the most up-regulated gene set, `TNFA_SIGNALING_VIA_NFKB`. In comparison, analyzing 442 randomly permuted gene lists with GSEAPreranked produced significant results in few experiments, less than 3% for any gene set in the Hallmarks collection.

**Figure 4:**
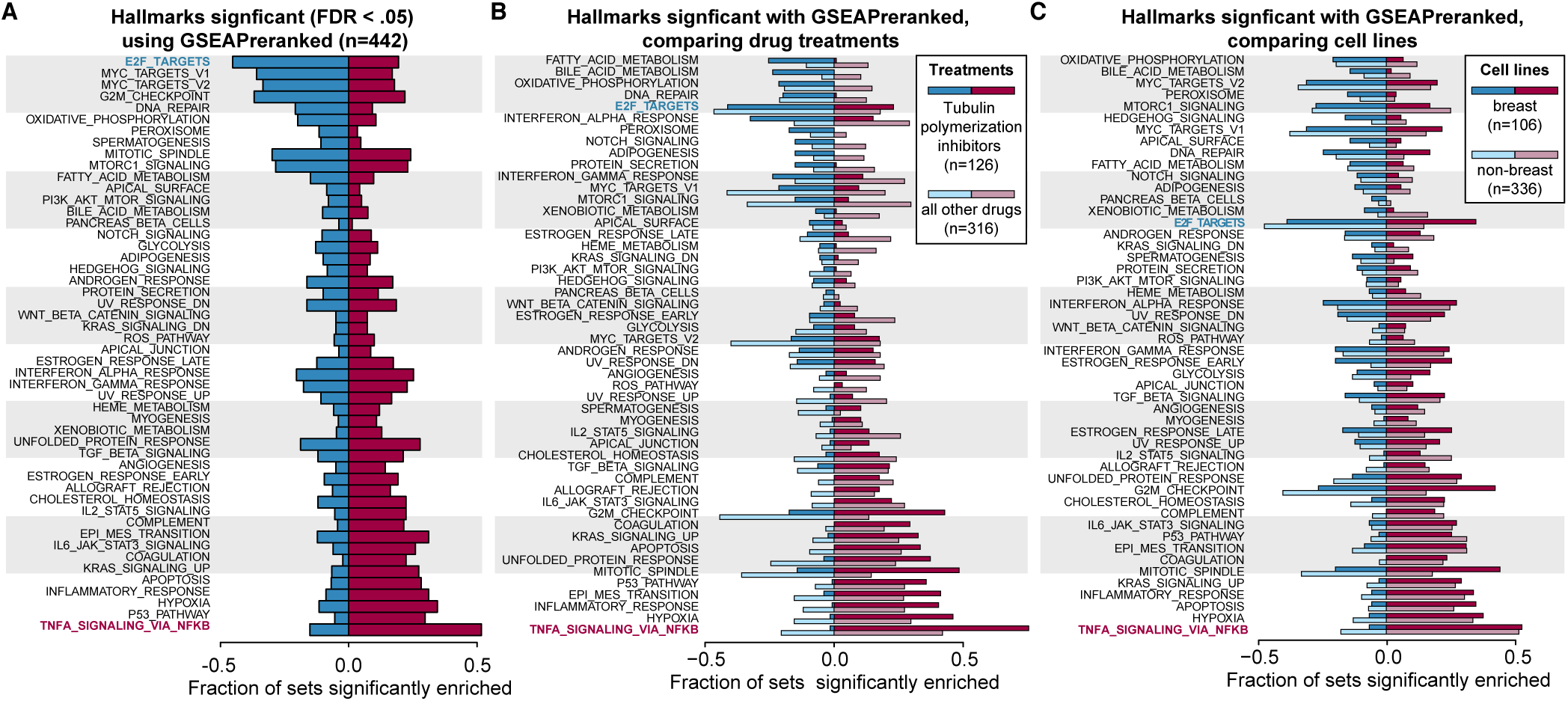
Commonly enriched gene sets across 442 small molecule gene expression experiments. (A) The gene sets in the Hallmarks collection [34] were tested against all 442 experiments using GSEAPreranked (competitive null hypothesis). Significant gene sets are defined as an FDR < 0.05. Gene sets are ranked by the difference in the fraction of experiments with significant positive and negative enrichment. The most frequently down-regulated pathway is `E2F_TARGETS` (blue text) and most commonly up-regulated pathway is `TNFA_SIGNALING_VIA_NFKB` (red text). (B) The fraction of positively and negatively enriched gene sets are shown for 126 experiments that tested response to eribulin or paclitaxel (dark bars), compared to 317 experiments that tested another compound (light bars). (C) The fraction of positively and negatively enriched gene sets within 107 experiments using breast cancer cell lines (dark bars), compared to 336 experiments that used non-breast cells (light bars).

To investigate the potential effects of the large number of experiments in our data set using breast cells or treating with tubulin polymerization inhibitors, we repeated our GSEAPreranked analysis including and excluding these experiments (Figure 4B). Comparing the GSEA results for 126 experiments using tubulin polymerization inhibitors to the remaining 317 experiments, we observed instances where certain gene sets increased in frequency of significance in experiments with the inhibitors and other gene sets increased under all other drugs. However, many of the general patterns shown in Figure 4A remain, demonstrating that the over-representation of tubulin polymerization inhibitors is not soley responsible for the results in Figure 4A. Similarly, we compared 107 experiments using breast cell lines to 336 experiments using cells from other tissues and, again, found gene sets such as `TNFA_SIGNALING_VIA_NFKB` were commonly significantly enriched regardless of experimental tissue type (Figure 4C).

### 3.3 Global adjustment of common patterns of gene set enrichment

Our meta-analysis of GSEA results across a compendium of 442 small molecule gene expression experiments highlighted common patterns of gene set enrichment. To complement this analysis, we next asked, which gene sets are uniquely enriched in a given experiment? We addressed this question by assuming the competitive null hypothesis as in GSEAPreranked, but adjusting the empirical null distribution used in the statistical test (Figure 1). The method we propose leverages a background set of experiments to define an informed null distribution, rather than creating one with completely random permutations. As the goal of this approach is to place a single experiment in the context of a background set of user-defined experiments, we call the method GSEA-InContext. Full details of the method are described in Section 2.3.

First, we compared the results produced by GSEAPreranked on the 442 experiments to the corresponding results produced by GSEA-InContext. We ran GSEA-InContext on each individual experiment using the background set of the 441 other experiments and the Hallmarks collection [34] (Figure 5). As expected, GSEA-InContext generally reduced the number of significantly reported gene sets per experiment. More specifically, the commonly enriched pathway `TNFA_SIGNALING_VIA_NFKB` was reduced from 53% up-regulated in GSEAPreranked to to 14% in GSEA-InContext. Similarly, the most down-regulated gene set `E2F_TARGETS` (42% enriched with GSEAPreranked) was enriched in only 19% of experiments using GSEA-InContext. Two Hallmark gene sets, `OXIDATIVE_PHOSPHORYLATION` and `PEROXISOME`, that are uncommon in the 442 experiments become enriched at a slightly higher frequency in GSEA-InContext compared to GSEAPreranked.

**Figure 5:**
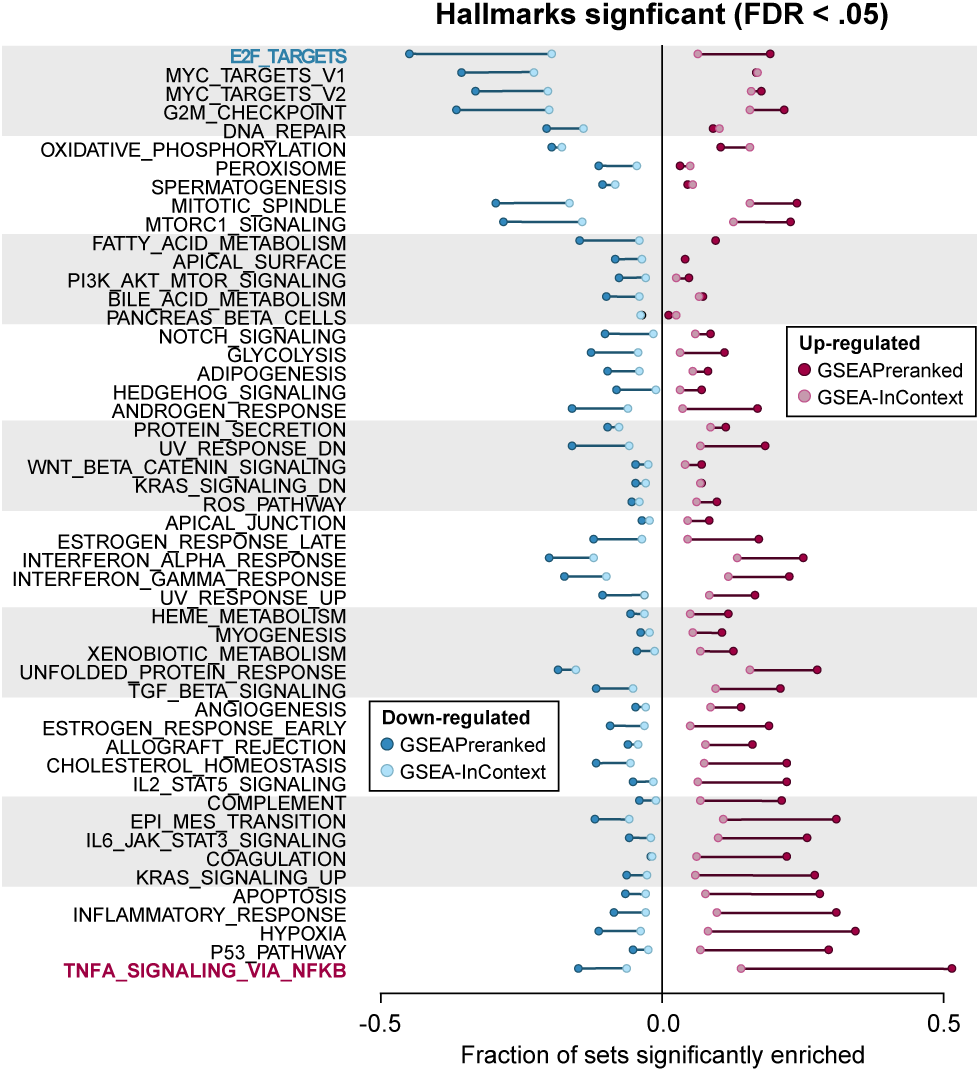
Adjusting for positively and negatively enriched pathways. The points represent the fraction of gene sets that are significantly up- or down-regulated (FDR < 0.05) across all 442 experiments in GSEAPreranked (dark red, dark blue) or GSEA-InContext (light red, light blue). The bars show the difference between the fraction of significantly enriched gene sets between the analyses.

To confirm that the GSEA-InContext method did not introduce any systematic biases, we ran GSEA-InContext on randomized rank lists for all of the 442 experiments. Similar to the findings using GSEAPreranked, we found that gene sets were significantly enriched in a small fraction of the random experiments, reaching a maximum of 3% of experiments significantly positively or negatively enriched.

### 3.4 Applications of GSEA-InContext

We demonstrate the application of GSEA-InContext using two biologically relevant examples. The first example illustrates that GSEA-InContext successfully removed non-specific gene set enrichment patterns in order to identify the on-target effects of a small molecule compound. The second example demonstrates how GSEA-InContext can be used to disentangle the effects of a single drug in cells treated with a drug combination.

#### 3.4.1 Re-scoring Notch pathway inhibition in a T-ALL cell line to down-weight nonspecific gene sets

Any small molecule drug will have direct (e.g., signaling) and indirect (e.g., stress) effects, whether it is due to drug promiscuity or the inherent interconnectedness of biological systems [25]. Thus, a perennial challenge in pharmacology is to functionally characterize the on- and off-target effects of a drug treatment. Accordingly, we demonstrate how GSEA-InContext can be used to identify gene sets that are specific to a small molecule treatment by selecting an appropriate background set of experiments. One well-represented tissue type in our compendium of 442 experiments is blood, in particular leukemia cell lines, which we stratified into the lymphoblastoid (n=44) and myeloid (n=48) lineages for this analysis. We selected a single experiment in which HBP-ALL cells treated with SAHM1, a Notch signaling inhibitor, were compared to cells treated with a vehicle control (GSE18198; [36]). Activation of Notch signaling has been associated with the development of T-cell acute lymphoblastic leukemia (T-ALL), and it has been shown that direct inhibition of Notch pathway members in tissue culture and mouse models decreases proliferation of T-ALL cells. Applying GSEAPreranked with the Hallmarks collection [34] to this experiment, we found 17 gene sets significantly enriched at an FDR < 0.05 (Figure 6A). Interestingly, while `NOTCH_SIGNALING` was down-regulated, it remained above the significance threshold (FDR = 0.097).

**Figure 6:**
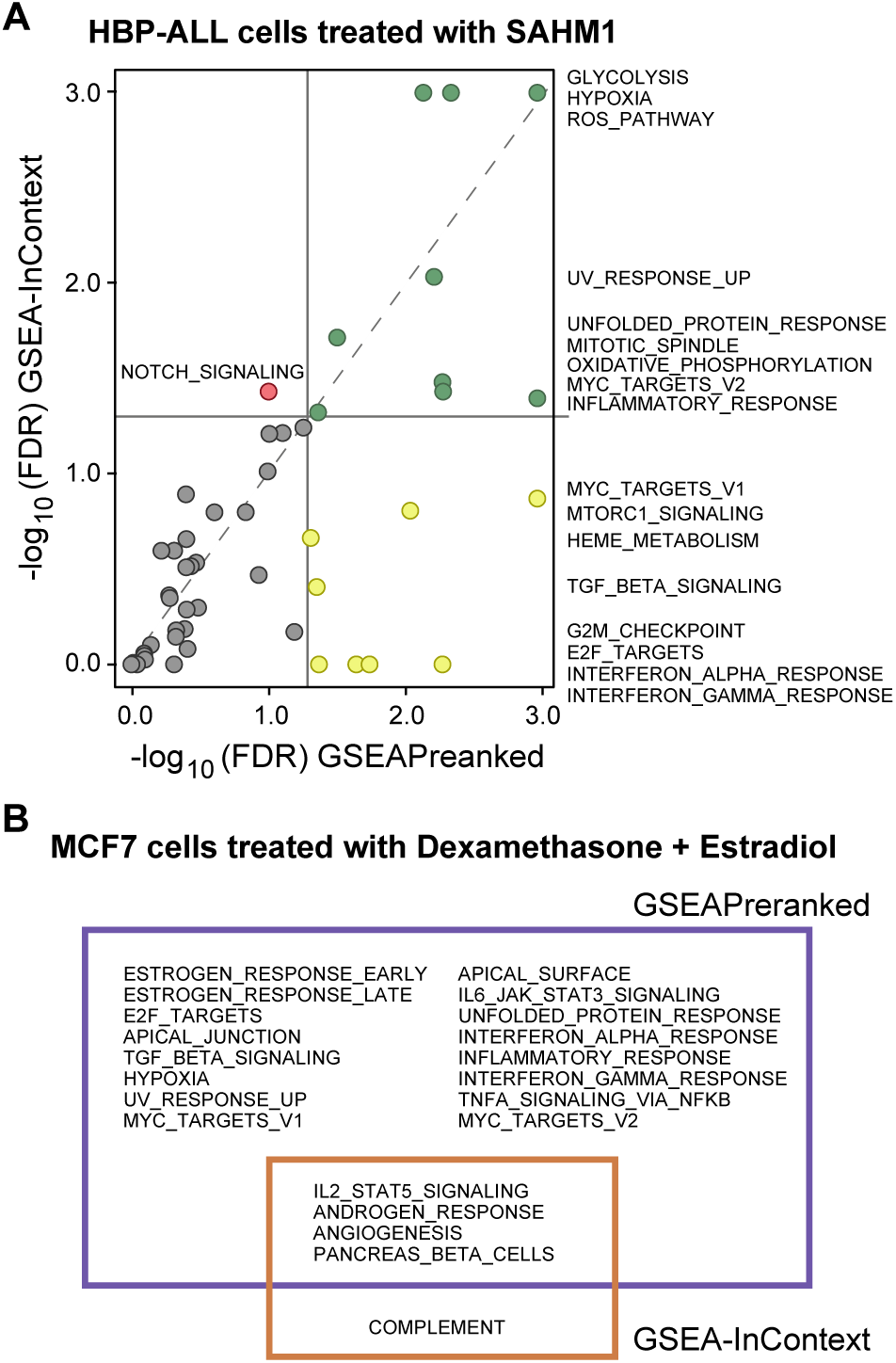
Two illustrative examples using GSEA-InContext. (A) GSEAPreranked was run on a ranked list of differentially expressed genes from T-cell acute lymphoblastic leukaemia cells (HBP-ALL) treated with SAHM1, a Notch pathway inhibitor. GSEA-InContext was run on the same experiment using a background set of 44 lymphoblastoid cell line experiments. The plot shows the −log_10_ FDR of each analysis, with the grey lines signifying FDR < 0.05. Yellow points represent gene sets significant only in GSEAPreranked; green points were signifcant in both analyses; the red point (`NOTCH_SIGNALING`) is only significant in GSEA-InContext; grey points fell below significance in both analyses. Gene set names are listed to the right of the plot. (B) GSEAPreranked significant results (purple) on an MCF7 breast cancer cell line treated with a combination of dexamethasone and estradiol. GSEA-InContext results (orange) on the same experiment as in (A) using a background set of 22 experiments in which MCF7 breast cancer cell lines were treated with estradiol only. The intersection and set differences between the two analyses are shown. All analyses used the genes sets in the Hallmarks collection [34].

We next ran the same experiment through GSEA-InContext, using a set of 44 lymphoblastoid experiments as the background set. Using the same Hallmarks collection, GSEA-InContext identified a total of 10 significantly enriched gene sets (FDR < 0.05). Notably, GSEA-InContext reported NOTCH_SIGNALING to be significantly down-regulated (FDR = 0.037) (Figure 6A), supporting the direct inhibition of the Notch signaling pathway by SAHM1 treatment. We confirmed that the direction of enrichment (positive/negative) for all gene sets was the same in both analyses.

We compared the results of GSEAPreranked to GSEA-InContext and used these patterns to help interpret the results. A gene set that was significant in GSEAPreranked but was raised above an FDR of 0.05 in GSEA-InContext was likely commonly enriched across the background experiments. Conversely, a gene set being significant in both GSEAPreranked and GSEA-InContext suggests that the set is uniquely enriched in the experiment being tested compared to the background experiments. We found gene sets that meet both criteria. Cell cycle related gene sets (`G2M_CHECKPOINT` and `E2F_TARGETS`) were significant in GSEAPreranked, but not in GSEA-InContext (Figure 6A), supporting the finding of Moellering, et al. that the SAHM1 inhibits cell proliferation [36]. Although down-regulation of cell cycle processes is a biologically relevant result that supports the authors experimental results, GSEA-InContext indicates that negative enrichment of cell cycle gene sets is a common response in lymphoblastoid cells treated with an array of drugs. This is supported by the fact that approximately 70% of the 44 background experiments showed enrichment of cell cycle related processes.

The most significantly down-regulated genes sets in GSEA-InContext are `HYPOXIA, GLYCOLYSIS, REACTIVE_OXYGEN_SPECIES_PATHWAY`. All three gene sets are also highly significant in GSEAPreranked, suggesting that these processes are uniquely significant when HBP-ALL cells are treated with SAHM1. The link between hypoxia and Notch signaling has been shown to play key roles in cell differentiation [23] and key cancer related processes of migration and invasion [40]. Hypoxia has long been know to play a key role in controlling glycolytic metabolism, particularly in cancer cells [35], and hypoxic conditions stimulate the production of reactive oxygen species [8]. The tight link between these process and their regulatory link with Notch suggests that Notch inhibition could be directly down-regulating key cancer progression processes, another potential positive effect of SAHM1 treatment.

Taken together, the GSEAPreranked and GSEA-InContext results provide a more complete picture of the pathways and processes that are differentially regulated in HBP-ALL cell treated with SAHM1. By placing enriched gene sets in context of a lymphoblastoid experimental background, we could identify both common and experiment-specific gene sets. In particular, GSEA-InContext identified NOTCH_SIGNALING as being significantly down-regulated, whereas GSEAPreranked did not (Figure 6A).

#### 3.4.2 Disentangling the effects of dexamethasone from estradiol response in breast cancer cell lines

For a second example, we sought to demonstrate how GSEA-InContext can be used to prioritize gene sets that are specific to a small molecule treatment by down-weighting gene sets that are enriched in the background set of experiments. In this case, we performed GSEAPreranked on an experiment in which MCF7 breast cancer cells were treated with estradiol, an estrogen receptor agonist, and dexamethasone, a corticosteroid (GSE79761) [48]. We then applied GSEA-InContext to this same experiment using a background set of 22 experiments in which MCF7 cells were treated with estradiol alone. By defining the background this way, we aimed to down-weight gene sets related to breast cancer cells or estradiol treatment while identifying gene sets that are more specifically related to dexamethasone treatment.

We compared the results for the Hallmarks collection [34] between each enrichment method (Figure 6B). Gene sets shown in the purple box in Figure 6B were significantly enriched using GSEAPreranked, but were not significantly enriched in GSEA-InContext. These gene sets represent pathways and processes that were commonly altered across the background experiments. In this group, we found gene sets that were expected to be enriched in MCF7 cells treated with estradiol, such as `ESTROGEN_RESPONSE_EARLY` and `ESTROGEN_RESPONSE_LATE`. Several gene sets that we previously identified as being significantly enriched across a wide variety of cell lines and drug treatments in our compendium (Figure 4), such as `E2F_TARGETS` and `TNFA_SIGNALING VIA_NFKB`, were also identified as significant by GSEAPreranked. In contrast, these gene sets were not significantly enriched in GSEA-InContext, demonstrating that these sets were successfully down-weighted to prioritize gene sets related to dexamethasone treatment while adjusting for the effects of estradiol.

The gene sets in the overlapping section between the purple and orange boxes were identified as significantly enriched in both GSEAPreranked and GSEA-InContext. We confirmed that the direction of enrichment (positive/negative) for these gene sets was the same in both methods. The four gene sets identified in both analyses were `ANGIOGENESIS, IL2_STAT5_SIGNALING, ANDROGEN_RESPONSE`, and `PANCREAS_BETA_CELLS`. Because these gene sets are also significant in the GSEA-InContext analysis, we expect the enrichment of these gene sets to be the result of the added dexamethasone treatment in these cells.

The link between dexamethasone and the androgen signaling pathway has been investigated in several studies. Dexamethasone is a glucocorticoid receptor (GR) agonist and GR shares several transcriptional targets with the androgen receptor (AR), including SGK1, MKP1, and DUSP1 [46]. Indeed, SGK1 is in the `ANDROGEN_RESPONSE` gene set. Dexamethasone has also been linked to IL2 signaling, which we see in the IL2_STAT5_SIGNALING gene set. The `ANGIOGENESIS` gene set is also negatively enriched in this experiment, supporting previous results showing that dexamethasone inhibits angiogenesis [51]. Finally, we note that `COMPLEMENT` is uniquely enriched in GSEA-InContext. Interestingly, dexamethosone has been shown to be a transcriptional regulator of components in the complement pathway [31]. While those results are in immune cells, this presents the potential research topic of dexamethosone regulation of complement in breast cells stimulated by estradiol.

Once again, we demonstrated that the GSEAPreranked and GSEA-InContext results taken together provide complementary perspectives into altered pathways and processes in this experiment. GSEAPreranked describes significant gene sets in our experiment compared to what would be expected at random, and by using GSEA-InContext to compare these enriched gene sets in the context of other MCF7 experiments treating with estradiol, we identified both common and experiment-specific gene sets.

## 4 Discussion

Extracting biological insights from the long lists of genes produced by differential expression experiments still remains a challenge. FCS methods, such as GSEA, are designed to aide in the interpretation of gene lists by identifying differentially up- and down-regulated pathways and processes. Although GSEA succeeds at summarizing the original list of genes into gene sets and identifying enrichment, the results are provided only in the context of the tested experiment. This is by design, but placing a single experiment in the context of a biologically relevant background can provide insight into the common and experiment-specific gene set patterns. In fact, common patterns of positively and negatively enriched gene sets can be observed across a variety of experimental conditions. We applied GSEAPreranked to 442 different experiments in which human cells were treated with small molecules and we identified gene sets that were commonly up- and down-regulated across a number of contexts (e.g. drugs and tissues).

The majority of drugs that we evaluated were inhibitors (most being cancer drugs). These small molecules are designed to inhibit the growth of cells. Consistent with what we expected, the gene sets representing cell cycle processes were the most down-regulated pathways, while gene sets associated with cellular damage and stress were commonly up-regulated. Interestingly, `TNFA_SIGNALING_VIA_NFKB` was significantly up-regulated in over 50% of the 442 experiments and NF-κB signaling downstream of TNFα has been shown to be pro-survival [38]. This suggests that inhibiting NF-κB signaling with the other small molecule could potentially be an effective drug combination treatment representing a common mechanism of drug synergy. This is one example of a testable hypothesis that can be generated from exploring commonly enriched gene sets.

Conversely, these common patterns motivate a new type of analysis: specifically, that researchers can place their own experimental results into a relevant context in order to identify uniquely enriched gene sets for their experiment compared to others. Accordingly, we introduced GSEA-InContext to perform such an analysis. By running GSEA-InContext on our compiled set of 442 expression experiments, we showed that the algorithm successfully down-weighted the gene sets such as `TNFA_SIGNALING_VIA_NFKB` that are commonly enriched in many experiments. Additionally, we applied GSEA-InContext to two example experiments, showing that in each case our method highlighted biological pathways relevant to the small molecule compound in each experiment.

The findings we present and the implementation of GSEA-InContext uses the competitive null hypothesis for statistically evaluating gene set enrichment. While the self-contained null hypothesis is preferred because it offers greater statistical power than the competitive null hypothesis [29, 19, 44], there are many instances when the self-contained null hypothesis cannot be used, particularly when the number of samples per condition are low. The majority of experiments that aim to test two conditions generate far less than seven samples per condition, which requires the competitive null hypothesis to be used for these experiments. Thus, while GSEA-InContext is not applicable using the self-contained hypothesis, it is is readily usable for the majority of gene expression experiments that require the use of the competitive null hypothesis.

For the purposes of this analysis, we focused our efforts on small molecule treatments of human cell lines. With over a million expression data sets currently in the GEO database [2], compiling a properly defined background set can be a daunting task, as each data set requires manual curation of the control and treatment groups. However, the 442 treatment-control comparisons that we compiled present a robust set of data to begin exploring common and experiment-specific gene set patterns. The results from GSEA-InContext will only become more robust as this background compendium is expanded to include other drugs and cell line experiments [30, 43]. Future work will also include compiling background sets to study other biological contexts and other organisms. Leveraging efforts such as CREEDS (CRowd Extracted Expression of Differential Signatures) will also rapidly expand the potential user-defined background sets [47]. Additionally, an area of future research will include studying platform-specific patterns to address any systematic biases that are introduced using hybridization technologies, such as GC content [49], and other technologies such as RNA-seq. Comparing results across platforms will help identify which commonly enriched gene sets can be attributed to technical differences between platforms and which patterns are robust across platforms and thus a true biological result.

We would like to close by stating that the goal of GSEA-InContext is not to replace the results of GSEA, but to complement the original implementation of GSEA [42, 37]. Comparing the results obtained from GSEA and the contextualized results from GSEA-InContext, we were able to gain insights into not only the pathway-level changes in an experiment, but also the common and experiment-specific patterns.

## Acknowledgements

We would like to thank Nicolle Witte for early contributions to the project.

